# Dynamics of the Human Antibody Repertoire following B-cell Depletion in Systemic Sclerosis

**DOI:** 10.1101/139758

**Authors:** Charles F. A. de Bourcy, Cornelia L. Dekker, Mark M. Davis, Mark R. Nicolls, Stephen R. Quake

**Author notes:** To whom correspondence should be addressed (e-mail), Clark Center E300, 318 Campus Drive, Stanford, CA 94305 (address). Note to Publishers: The authors are in the process of deciding to which journal to submit the present manuscript. Interested editors should contact Stephen R. Quake.

## Abstract

Systemic sclerosis with pulmonary arterial hypertension (SSc-PAH) is a debilitating and frequently lethal disease of unknown cause lacking effective treatment options. Lymphocyte anomalies and autoantibodies observed in systemic sclerosis have suggested an autoimmune character. Here we study the clonal structure of the B-cell repertoire in SSc-PAH using immunoglobulin heavy-chain sequencing before and after B-cell depletion. We found SSc-PAH to be associated with anomalies in B-cell development, namely altered VDJ rearrangement frequencies (reduced IGHV2-5 segment usage) and an increased somatic mutation-fixation probability in expanded B-cell lineages. SSc-PAH was also characterized by anomalies in B-cell homeostasis, namely an expanded IgD^+^ proportion with reduced mutation loads and an expanded proportion of highly antibody-secreting cells. Disease signatures pertaining to IGHV2-5 segment usage, IgD proportions and mutation loads were temporarily reversed after B-cell depletion. Analyzing the time course of B-cell depletion, we find that the kinetics of naïve replenishment are predictable from baseline measurements alone, that release of plasma cells into the periphery can precede naïve replenishment and that modes of B-cell receptor diversity are highly elastic. Our findings shed light on the humoral immune basis of SSc-PAH and provide insights into the effect of B-cell depletion on the antibody repertoire.

**Abbreviations:**

SSc-PAH: Systemic sclerosis with pulmonary arterial hypertension
IGH: immunoglobulin heavy-chain
BCR: B-cell receptor
CDR3: complementarity-determining region 3
Ig: immunoglobulin
AA: amino acid

## Introduction

Systemic sclerosis (SSc), or scleroderma, is a rare chronic autoimmune disease leading to tissue fibrosis and vasculopathy (1). Indications of a dysregulated immune system include reduced lymphocyte counts (2), T-cell anomalies (3) and B-cell anomalies, in particular the presence of autoantibodies (4), increased CD19 expression (5), increased naïve B-cell proportions and diminished memory B-cell proportions (5). A complication with particularly poor prognosis is pulmonary arterial hypertension (PAH) (6), which can arise from SSc-associated interstitial lung disease (ILD), but also directly from SSc without significant ILD; the latter case, which we will refer to as SSc-PAH, is the object of the present study. Current treatment typically focuses on pulmonary vasodilation and is not curative (7). Immunotherapy with the B-cell depleting agent rituximab, a monoclonal antibody against CD20 (8, 9), has shown promise as a potential treatment for some forms of SSc, improving disability scores (10) and lung function (11), but has not previously been assessed in controlled multicenter clinical trials for SSc-PAH. The B-cell basis of SSc has previously been studied (12) at the level of subset sizes (5), cytokine levels (13), response regulators (5, 14) and autoantibody specificities (15, 16), but not at the level of sequence-based B-cell phylogeny. Reconstitution of the immunoglobulin heavy-chain repertoire following rituximab administration has been studied before (17), but sample size was small (2 participants) and sequencing depth was low (on the order of 680 sequences).

Here, we elucidate signatures of SSc-PAH and dynamics of B-cell replenishment after depletion using massively parallel sequencing of immunoglobulin heavy-chain (IGH) transcripts found in peripheral-blood mononuclear cells (“antibody repertoire sequencing”), which provides a wealth of information on the processes of VDJ recombination, clonal expansion, somatic hypermutation and isotype-switching that shape the B-cell repertoire (18, 19). We longitudinally studied 11 participants from an ongoing randomized, double-blind, placebo-controlled phase II multicenter trial of rituximab for the treatment of SSc-PAH, sequencing 85 samples and obtaining > 4 million IGH sequences. Infusions of 1000 mg of rituximab or a placebo were given at weeks 0 and 2; blood was sampled at baseline (pre-infusion), at weeks 2 and 4 relative to the first infusion, and roughly in 12-week increments after that (Fig. 1A). All SSc-PAH participants were females between the ages of 48 and 73, inclusive; healthy baseline controls were 15 females between the ages of 50 and 75, inclusive (Suppl. Table S1).

**Fig. 1.**
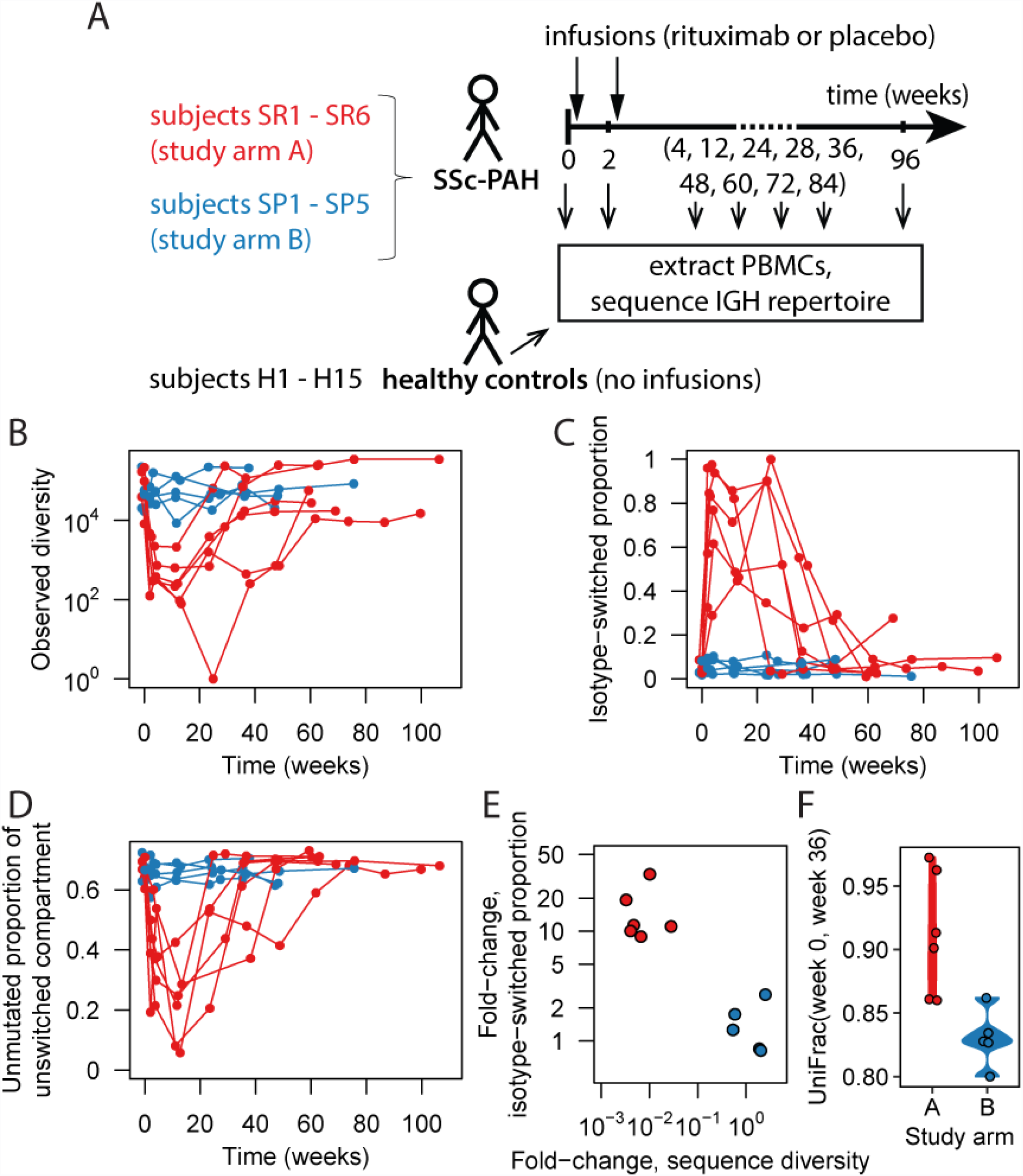
Study design and comparison of study groups across time. (**A**) Study design. PBMCs, peripheral blood mononuclear cells. Whether infusions were of rituximab or placebo depended on the study arm. (Note that not all the indicated time points were available for all participants.) (**B**) Number of distinct IGH sequences observed as a function of time, colored by study arm. (**C**) Fraction of observed sequences that had the IgA or IgG isotype as a function of time, colored by study arm. (**D**) Fraction of IgD or IgM sequences that displayed zero somatic mutations as a function of time, colored by study arm. (**E**) Summary of the signatures identified in (B) and (C): fold-change in isotype-switched proportion from baseline to depletion versus fold-change in sequence diversity from baseline to depletion. Here “depletion” refers to the week 4 and week 12 visits; diversities and isotype-switched proportions were averaged across those two time points. (**F**) Phylogenetic distance between each participant’s baseline repertoire and her repertoire at the week 36 visit. Week 36 was chosen as the common second time point because all but two arm-A recipients had recovered more than 10,000 sequences by then and times beyond week 36 had not been sampled for all participants.

## Results

### Identification of a study group undergoing B-cell depletion

Since the clinical trial providing study samples is ongoing, it is blinded as to whether study arm A or B corresponded to the rituximab group as opposed to placebo (Fig. 1A). Here, we show that the characteristics of group A are consistent with CD20^+^ B-cell depletion. Indeed, group A shows a transient reduction in overall B-cell receptor (BCR) diversity (Fig. 1B) and the vast majority of B-cells remaining during the depletion period have undergone isotype switching (Fig. 1C) or acquired one or more mutations (Fig. 1D), consistent with the hypothesis that they are CD20-B-cells in the terminal stages of differentiation (plasmablasts and plasma cells (20)). B-cell depletion was observed to be transient, ended by repopulation with new naïve cells (Fig. 1B,C,D). Summarizing the above depletion signatures as fold-changes between baseline and early post-infusion time points, it is clear that subjects cluster into an unaffected and an affected group (Fig. 1E). In order to compare the initial B-cell repertoire to the reconstituted repertoire after depletion, we applied the phylogenetic distance metric UniFrac (21) whose use on immune repertoire data was recently demonstrated (22). As expected, we found that group A displayed greater pre-to-post-infusion distances than group B (Fig. 1F), consistent with repertoire reconstitution by a fresh set of naïve B-cells not clonally related to the baseline population. These observations establish group A as a cohort that can be used to study B-cell depletion.

### B-cell signatures of SSc-PAH

The hypervariable complementarity-determining region 3 (CDR3) of antibodies is a key determinant of antigen specificity (23). Antibody repertoire analysis has been used to search for convergent CDR3 sequences characterizing the immune response to certain infectious agents, notably dengue virus (24). To investigate which antibodies may be responsible for autoimmunity in systemic sclerosis, we tested all CDR3 amino acid sequences identified in this study, together with their one-mismatch derivatives, for enrichment in the SSc-PAH cohort versus the healthy cohort (Fisher Exact Tests with Benjamini-Hochberg correction). No sequences displayed a statistically significant association with SSc-PAH (Suppl. Fig. S1A,B). This negative result may be due to limited statistical power or undersampling of the repertoire, but also to biological idiosyncrasy: the relevant autoantigen(s) may differ from participant to participant and antibodies binding any given autoantigen may be encoded by distinct amino acid sequences.

We may nevertheless gain insight into the disease by studying the global structure of the antibody repertoire rather than individual antibody sequences. SSc-PAH participants tended to display slightly lower overall BCR diversity (Suppl. Fig. S1C), a higher proportion of IgD diversity and lower mean mutation loads in IgD than healthy subjects (Fig. 2A), consistent with previous reports of reduced lymphocyte counts (2) and expanded naïve B-cell proportions (5). This finding constitutes evidence for altered B-cell homeostatic mechanisms.

**Fig. 2.**
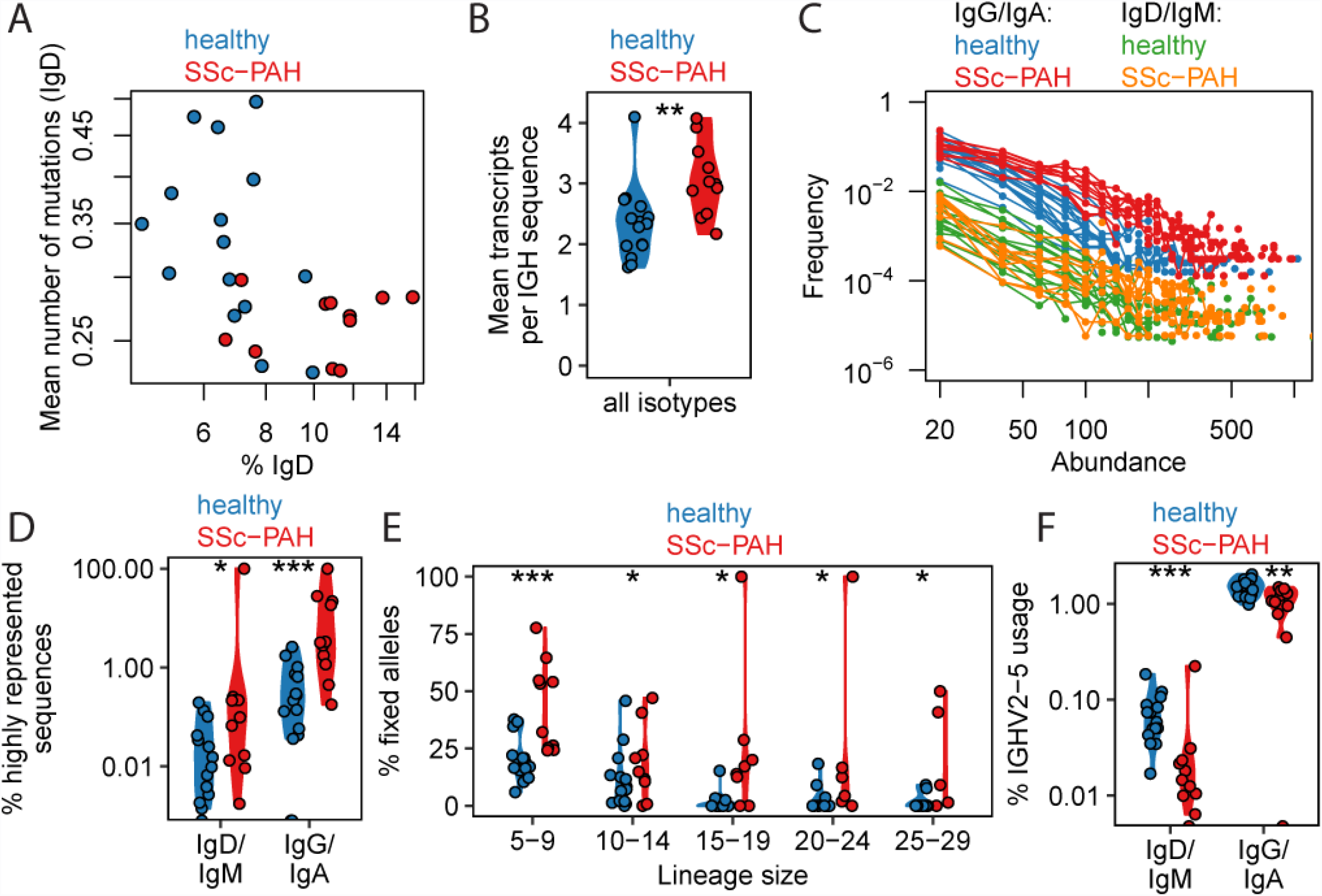
Signatures of SSc-PAH compared to healthy Ig repertoires at baseline. (**A**) IgD proportion of the IGH sequence repertoire and average mutation load in IgD. (**B**) Average number of transcripts recorded per distinct IGH sequence (i.e. sum total of all transcripts across all observed IGH sequences divided by the number of observed IGH sequences). Each point corresponds to one study subject. Significance codes: “*” 0.01 ≤ p < 0.05, “**” 0.001 ≤ p < 0.01, “***” 0.0001 ≤ p < 0.001. (**C**) Distribution of transcript abundances, separated by isotype compartment. Each curve corresponds to one study subject. Here “abundance” is the number of transcript copies associated with a sequence and “frequency” refers to the number of distinct sequences that display a given abundance value. (**D**) Percentage of IGH sequences that had a relative transcript abundance of at least 0.05% of the total repertoire, separated by isotype compartment. Points straddling the lower edge of the panel correspond to the value 0, which cannot be rendered on the logarithmic scale. (Note: when the single high value in SSc-PAH IgD/IgM is removed (subject SR3), the p-value for the IgD/IgM comparison changes from 0.026 to 0.049 (< 0.05); there are 2 points at zero for healthy IgM/IgD). (**E**) Proportion of somatic mutations that were considered to have become fixed in their B-cell lineage, i.e. present in at least 80% of sequences in that lineage. The proportion of fixed mutations was calculated separately for each lineage, then averaged across all lineages in the relevant size bin. Here lineage size refers to the number of distinct IGH sequences in the lineage. (Note: the two highest points for the “15-19” bin, the highest point for the “20-24” bin and the two highest points for the “25-29” bin correspond to 5 distinct SSc-PAH subjects, namely SP5, SP4, SP2, SR4, SR2). (**F**) Percentage of lineages that used the IGHV2-5 segment for each isotype compartment. Points straddling the lower edge of the panel correspond to the value 0, which cannot be rendered on the logarithmic scale. (Note: when the single high point for SSc-PAH IgD/IgM is removed (subject SR3), the p-value for the IgD/IgM comparison changes from 5.1×10^−4^ to 7.3×10^−6^. When the single point at zero in SSc-PAH IgG/IgA is removed (subject SP2), the p-value for the IgG/IgA comparison changes from 4.4×10^−3^ to 9.6×10^−3^.)

It has also been reported that memory B-cells are reduced in number but activated in scleroderma (5). Here, we are able to obtain further insight into the anomalies of the antigen-experienced compartment. We found that the average expression of distinct BCRs was higher in SSc-PAH (Fig. 2B), that this increase in average expression was due to an accumulation of BCRs at the highest observed expression levels (Fig. 2C), and that the effect was mostly driven by the isotype-switched (i.e. antigen-experienced) compartment (Fig. 2D). The effect persisted when abundances were defined in a relative rather than absolute fashion, i.e. as a proportion of the total repertoire abundances, eliminating any undesired potential systematic variations in sequencing depth (Fig. 2D). IGH sequences observed at high abundance are likely to correspond to plasmablasts, so the excess diversity of high-abundance BCRs suggests B-cells differentiate more frequently into plasmablasts in SSc-PAH participants than in healthy subjects.

In order to further elucidate the importance of selection processes acting on proliferating B-cell lineages, we measured what fraction of somatic mutations had become fixed in each lineage, defining a fixed mutation as one that was present in at least 80% of the lineage’s sequences. (A lineage was defined as a set of sequences with the same V-segment, same J-segment, same CDR3 length and 90% CDR3 sequence similarity in single-linkage clustering.) To eliminate artifacts due to differences in lineage sizes, the analysis was carried out separately for different ranges of lineage sizes. Germline alleles distinct from the reference could be misinterpreted as fixed somatic mutations, but no convincing evidence for the presence of novel germline alleles was found using TIgGER (25) on the present data set. We found that the SSc-PAH cohort on average had a larger fraction of fixed mutations than the healthy cohort across lineage sizes of up to 30 sequences (larger lineages were not consistently found in subjects) (Fig. 2E). This finding suggests that SSc-PAH lineages have on average undergone more selective sweeps, i.e. have been subject to sustained affinity maturation in response to antigen. One can speculate that the antigens in question are self-antigens, perhaps triggered by molecular mimicry (26) and not necessarily the same for each patient.

To assess whether VDJ rearrangement probabilities or post-rearrangement tolerance checkpoints (27) may be affected in SSc-PAH, we tested for differential expression of V-segments, which form the most diverse set of germline segments compared to D and J. In order to focus on original VDJ recombination events while removing the effects of clonal expansion and transcript abundance, V-segment counts were defined as a number of lineages (rather than number of sequences or number of transcripts). We found that IGHV2-5 was used at a lower level in SSc-PAH participants (Suppl. Fig. 1D, Fig. 2F). Note that IGHV2-5 usage was higher for the isotype-switched than for the unswitched compartment (Fig. 2F), but differential expression between disease and healthy cohorts was observed separately in both compartments, so the effect cannot be solely a consequence of the SSc-PAH-related expansion of the unswitched compartment (Fig. 2A). Differential expression was particularly pronounced in the unswitched compartment (Fig. 2F).

Having identified these B-cell signatures of SSc-PAH, we can ask how they are affected in subjects undergoing B-cell depletion (study arm A). Interestingly, in study arm A, the SSc-PAH-related excess of IgD diversity and deficit in mean IgD mutation load were transiently reversed (Fig. 3A,B), as was the deficit in IGHV2-5 usage (Fig. 3C). By contrast, B-cell depletion did not have a clear effect on the fraction of fixed mutations (Fig. 3D).

**Fig. 3.**
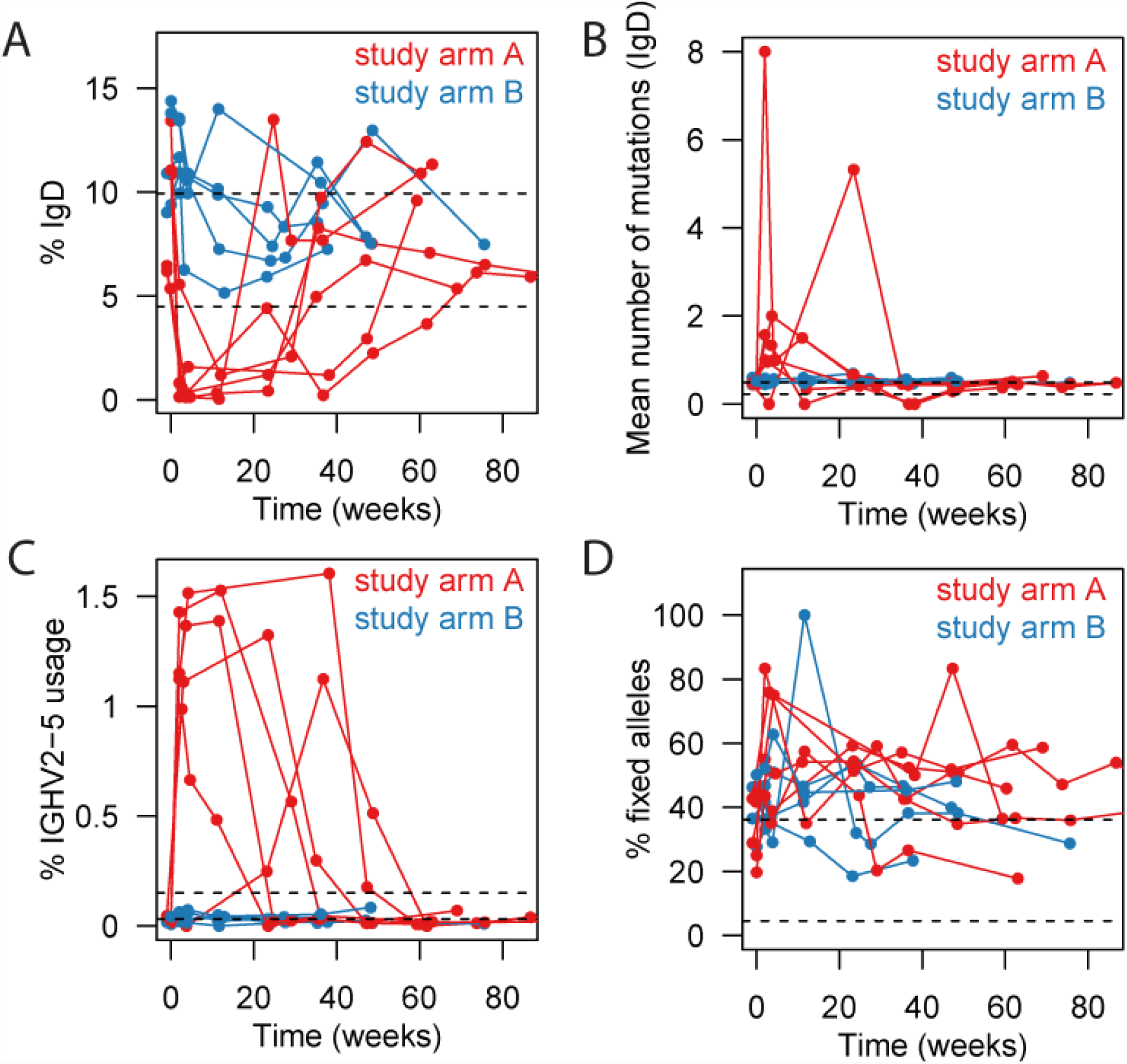
Time courses of SSc-PAH signatures. Dashed black lines indicate the range of values observed in healthy subjects at baseline. Note that the data displayed for study arms A and B in the present figure (longitudinal experiment) was acquired in a separate batch from the data on healthy subjects (cross-sectional experiment, already described in Fig. 2) and so may not be directly comparable to the healthy data; we nevertheless include the dashed lines as a guide. (**A**) IgD proportion. (**B**) IgD mutation loads. (**C**) Percentage of lineages using the IGHV2-5 segment (all isotypes combined). (**D**) Proportion of somatic mutations observed in a lineage that were considered “fixed”, i.e. present in at least 80% of the lineage’s sequences. The proportion of fixed mutations was calculated separately for each lineage containing between 5 and 29 distinct sequences, then averaged across all those lineages.

### Kinetics of repertoire reconstitution

Since we have inferred that the peripheral naïve repertoire is almost entirely wiped out in study group A (Fig. 1B,C,D,E), we can use that group’s longitudinal repertoire data to deduce various rates governing homeostasis of the humoral immune system. Using the number of IgM sequences with zero somatic mutations as a proxy for the naïve diversity of the repertoire, we can express the rate of change of naïve diversity as:

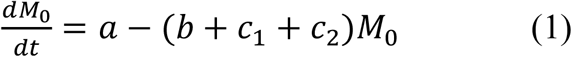

where *M_x_* denotes the number of IgM sequences with *x* mutations, *a* is the generation rate of naÏve sequences entering the periphery from the bone marrow, *b* is a mutation probability per unit time, *c*_1_ is a class-switch probability per unit time, and *c*_2_ is a probability of apoptosis per unit time. Sequences that exit the M_0_ diversity by acquiring a mutation will increase the M_1_ diversity:

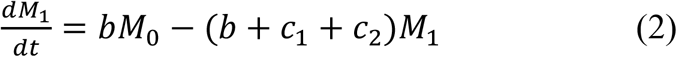

Cells can leave the IgM compartment by either undergoing apoptosis (rate *c*_2_) or by switching isotype (predominantly to IgA or IgG, rate *c*_1_). In the latter case they increase the isotype-switched diversity:

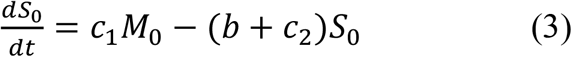

where *S_x_* denotes the number of IgA/IgG sequences with *x* mutations. In equations (1) and (2), we have made the simplifying assumption that the “exit” rates *b*, *c*_1_ and *c*_2_ applicable to the M_0_ diversity are the same for the M_1_ diversity; similarly, in equation (3), we have assumed that the rates *b* and *c*_2_ governing exits from the S_0_ diversity have the same values as in equations (1) and (2). Indeed, for a first approximation, it seems reasonable to assume that the fraction of a compartment that undergoes apoptosis/mutation/class-switching is roughly independent of the identity of the compartment, particularly if we restrict ourselves to lowly-mutated compartments that likely have a smaller proportion of long-lived memory/plasma cells than more highly mutated compartments.

Given that assessment of diversity using repertoire sequencing is a reliable measurement (Suppl. Fig. S2A,B), we can determine *a* and *d* ≡ (*b* + *c*_1_ + *c*_2_) from the B-cell replenishment time course using equation (1), starting at the onset of replenishment (after the more or less extended near-total depletion period). Fitting the linear model for all groups simultaneously as described in Suppl. Fig. S2C (after correcting observed sequence numbers for unseen sequences using the Chao1 estimator (28) suggested that coefficient *d* varied little between participants whereas coefficient *a* varied significantly, so we fitted the data again using a reduced model with a universal coefficient *d* for all participants and coefficients *a* that were allowed to vary from participant to participant (fits shown in Fig. 4A). As expected for a steady-state diversity controlled by these generation and exit rates, the diversity to which the M_0_ compartment returns after depletion is commensurate with its initial diversity (Suppl. Fig. S2D).

**Fig. 4.**
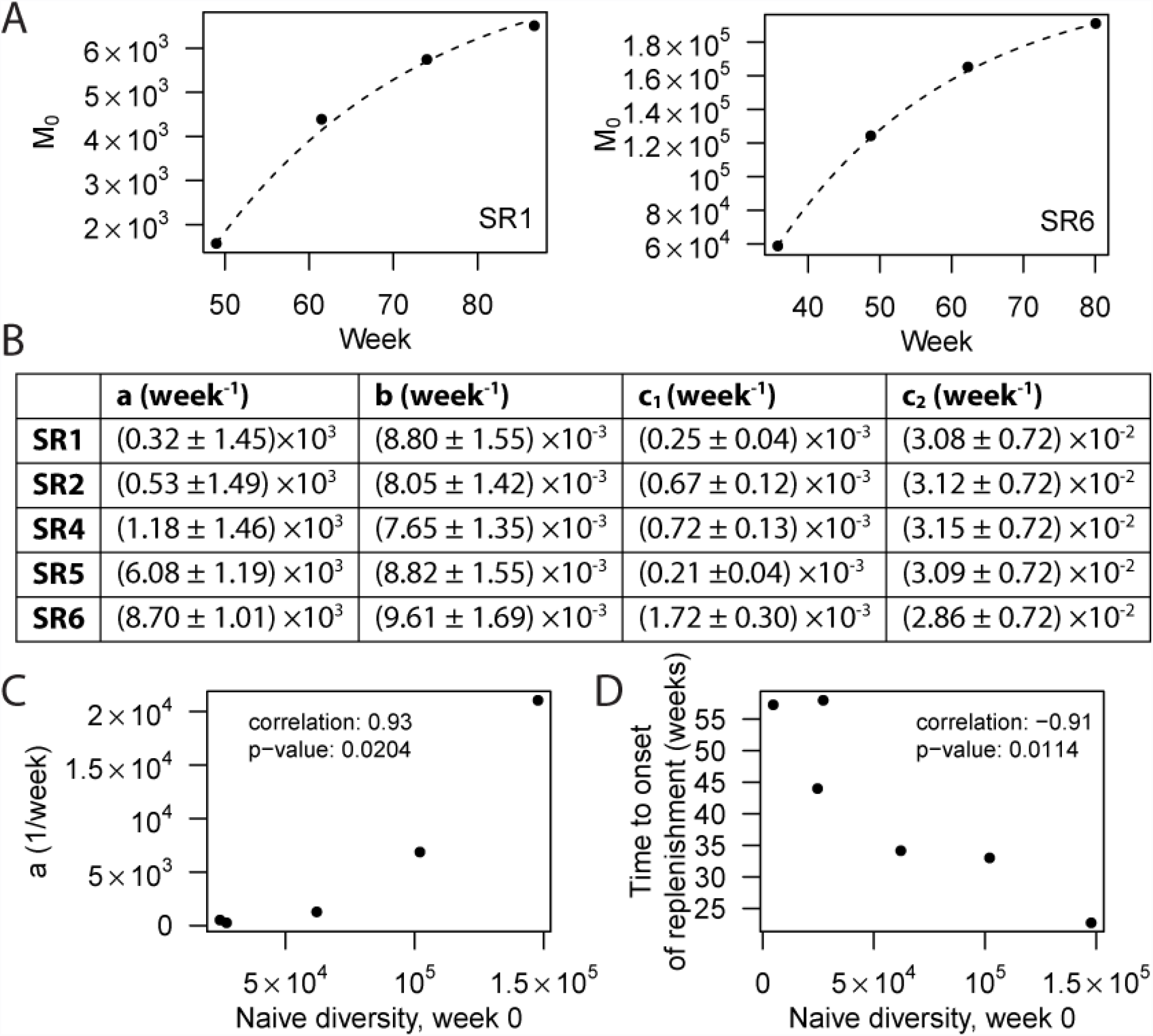
Quantitative modeling of repertoire replenishment rates. (**A**) Fit for the naïve diversity M_0_ (number of non-mutated IgM sequences, Chao1 estimate) as a function of time, following onset of replenishment. Only the two participants (SR1, SR6) with the largest numbers of corresponding time points are shown. (**B**) Fit parameters describing sequence generation (a), mutation (b), class-switch (c1) and apoptosis (c2) rates. SR1-SR6 are labels for individual subjects. Participant SR3 is omitted because an insufficient number of time points after onset of depletion were available. (**C**) Baseline naïve diversity versus naïve generation rate. (**D**) Baseline naïve diversity versus time to onset of replenishment (defined as the interval between second visit and the earliest visit exhibiting more than 5000 distinct IGH sequences).

In steady-state, i.e. in the baseline homeostatic regime, the time derivatives in equations (2) and (3) are zero, so we can deduce 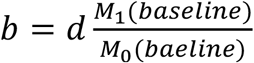 (4) and 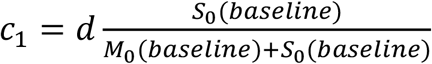 (5), followed by *c*_2_ = *d* − *b* − *c*_1_ (Fig. 4B). Note that the steady-state form of equation (2) can be generalized to higher mutation loads: 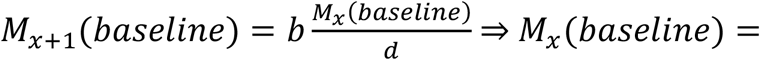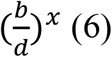, meaning the distribution of mutation loads in the IgM compartment should be an exponential decay, which is roughly obeyed in our data (Suppl. Fig. S2E; note that the isotype-switched compartment displayed in Suppl. Fig. S2F follows a contrasting distribution peaked at a non-zero mutation load). Since the data shows a hint that there may be a slight excess of highly-mutated IgM sequences compared to the exponential law, which may be attributable to long-lived IgM memory cells and plasmablasts that do not follow our simple system of equations above, we decided to determine *b* based on only M_0_ and M_1_ using equation (4) rather than fitting the entire distribution Mx using equation (6). The resulting sequence generation, mutation, class-switch and apoptosis rates (Fig. 4B) show that apoptosis due to the absence of activation by any cognate antigen is the most important factor limiting the naïve diversity; once a cell gets activated by a cognate antigen, a mutation event is more likely to occur first than a class-switching event (c2 > b > c1).

If one wanted to predict the time course of naïve B-cell replenishment after the first depleted time point, one would need three pieces: the time to onset of replenishment, the generation rate *a*, and the exit rate *d*. Now, *d* appeared to be near-“universal” between participants, *a* was found to be strongly correlated with the baseline naïve diversity (Fig. 4C), and the time to onset of replenishment was found to be inversely correlated with baseline diversity (Fig. 4D). These observations suggest the intriguing possibility of predicting depletion/replenishment time courses from baseline measurements alone.

### Similarities and perturbations of the reconstituted repertoire compared to baseline

We sought to assess which features of the BCR repertoire are elastic or plastic under B-cell depletion. We found that there is a clear stereotypy to the repertoire that is recovered after depletion: isotype usages, VDJ combination usages and mutation-load histograms show a marked similarity to their baseline profiles that is only temporarily lost after depletion (Fig. 5A). The mutation load profiles show that high mutation loads do not necessitate a lifetime of pathogen-induced affinity maturation, but are recovered within months – so any subsequent improved immune responses are presumably not generated by B-cell lineages acquiring more and more mutations ad infinitum, but rather by adding breadth to the phylogenetic tree. Frequencies of VDJ segment combinations (defined here as the number of distinct IGH sequences with a given combination) exhibited a similarity between participants at baseline that was lost in depletion but recovered at the monitoring endpoint (Suppl. Fig. S3A). The set of amino acid (AA) changes in the repertoire also showed a greater degree of inter-participant similarity when repertoires were full (baseline or endpoint) rather than depleted (Suppl. Fig. S3B). This observation suggests intrinsic biases of the somatic hypermutation machinery that are conserved between participants (29), contrasted with participant-by-participant idiosyncrasy of the mutations exhibited by the (presumably highly pathogen-specific) plasmablasts that remain during depletion. While VDJ usages, isotype usages and mutation histograms were highly elastic – demonstrating that mechanisms of diversity generation are intact after B-cell depletion – only a fraction of AA changes present at baseline were observed to return by the endpoint (Fig. 5A, “specific mutations”).

**Fig. 5.**
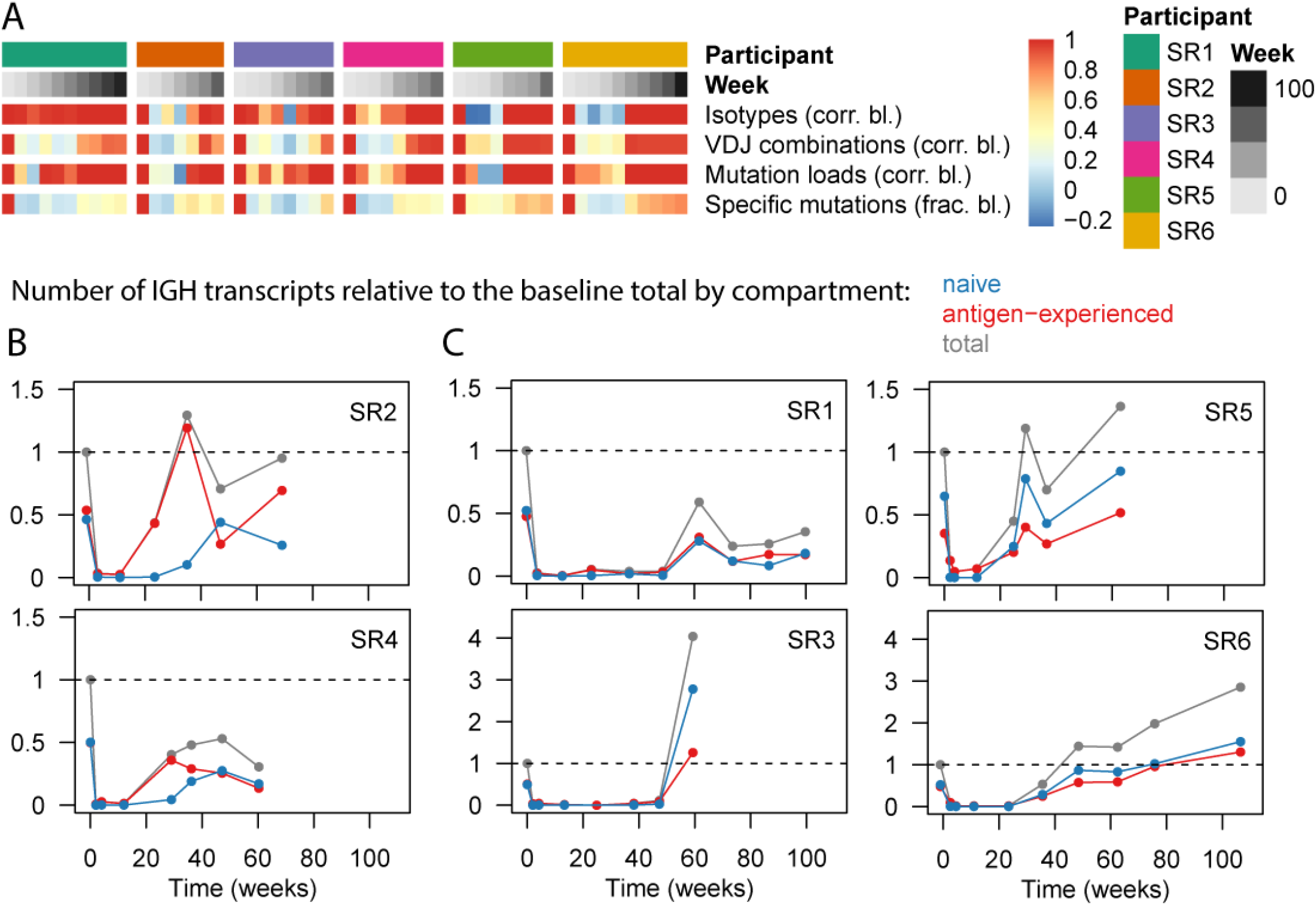
Assessment of repertoire elasticity and compartment replenishment after depletion. (**A**) Similarity of various repertoire characteristics to their baseline state on a per-participant basis. The top two rows identify participant and time point; the other rows display data as a heat map for each of the indicated variables, compared to baseline using either a correlation coefficient or a fraction as indicated. Naturally, for each participant, the baseline value is 1; values then tend to drop during repertoire depletion and gradually increase again during replenishment. For isotype usage profiles (resolved by subisotype) and VDJ usage profiles, the value shown at each time point is the correlation to the baseline profile. A repertoire’s “mutation load profile” was defined as the number of nucleotide sequences having each mutation load from 0 to 10 (nucleotide) mutations; we display the correlation of such mutation load histograms to the baseline histogram. “Specific mutations” refers to the set of distinct V-segment amino acid mutations observed in a repertoire (each defined by the identity of the V segment, the position being mutated, and the identity of the observed residue). We displayed the fraction of baseline mutations that were observed at each post-baseline time point in order to indicate to what extent specific mutations observed at baseline return after B-cell depletion. (**B,C**) Number of transcripts observed as a function of time, separated by compartment (naïve or antigen-experienced), relative to the total number of transcripts (from either compartment) observed at baseline. Notice that in (B) the antigen-experienced fraction increases well before the naïve fraction while in (C) it does not. Sequences were considered naïve if they were IgD or IgM and non-mutated; antigen-experienced if they had at least one mutation or were class-switched.

Next, we disentangled the B-cell populations that replenish the repertoire. First, we contrast antigen-experienced cells with naïve cells (Fig. 5B,C), focusing on their contributions to the total transcript abundances, which should be a proxy for rough protein abundances. We observe two distinct phenotypes: in certain participants (SR2 and SR4), the first post-depletion repertoire consists almost entirely of antigen-experienced transcripts (Fig. 5B), whereas in most participants (SR1, SR3, SR5, SR6) the naïve and antigen-experienced compartments recover simultaneously (Fig. 5C). The latter phenotype is what one might expect in a model in which new naïve B-cells enter the periphery from the bone marrow, soon encounter antigen, and get activated. The former phenotype suggests a scenario in which an early surge in plasmablasts or plasma cells palliates B-cell depletion until replenishment of naïve cells begins. Since plasmablasts live only a few days before dying or terminally differentiating into long-lived plasma cells in the bone marrow (30), this phenotype must be explained by long-lived plasma cells being released from the bone marrow to populate the blood. The early increase in antigen-experienced transcripts in subjects SR2 and SR4 corresponded to increases in the IgA1, IgA2 and IgG2 isotypes beyond baseline levels (Fig. 6A); in subject SR2 there was also a marked increase in IGH sequences with high somatic mutation loads (range 6-9 mutations in the sequenced region, Fig. 6B).

**Fig. 6.**
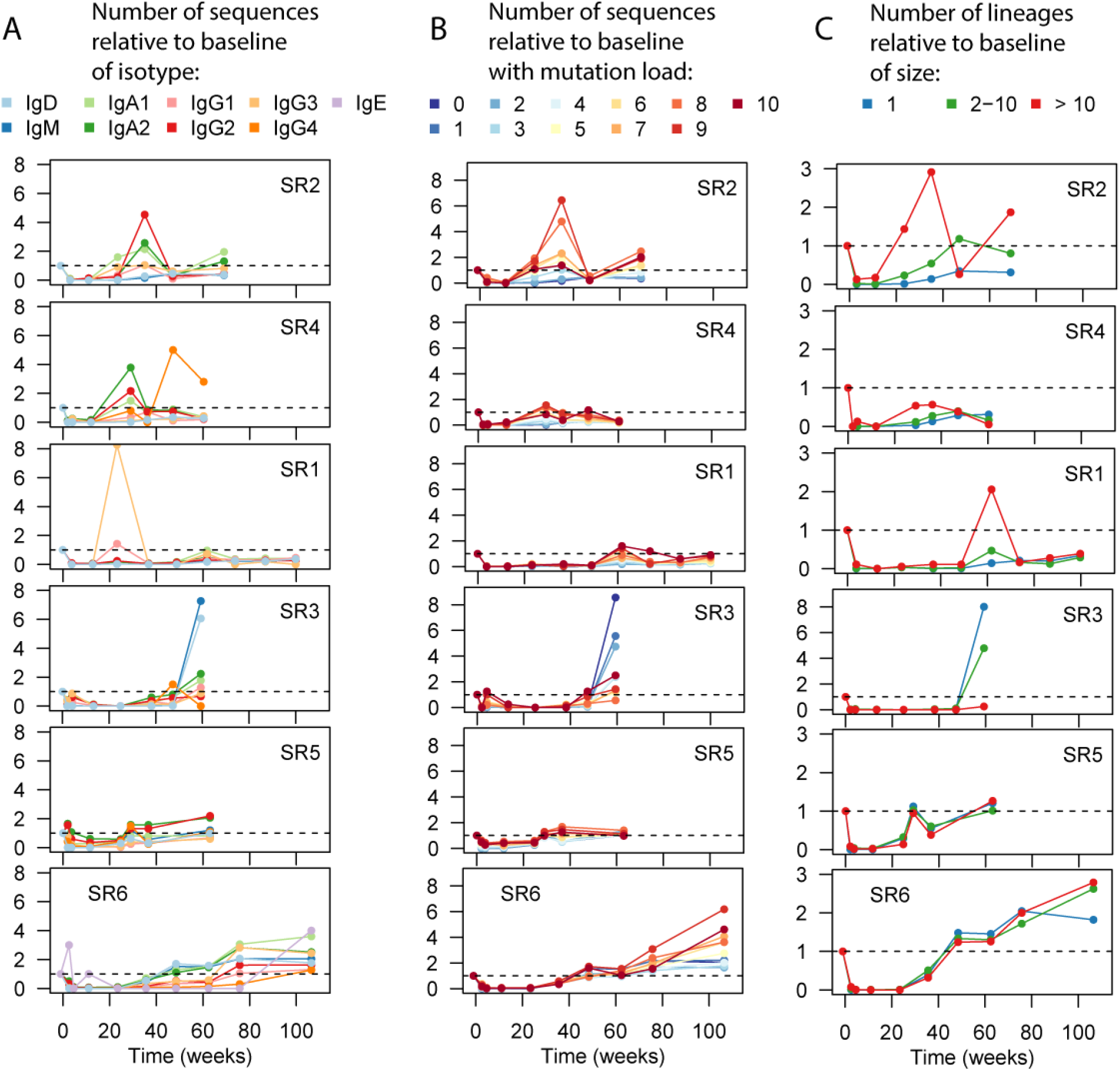
Comparison of repertoire distributions to baseline. (**A**) Frequency of sequences with each subisotype relative to baseline. (**B**) Frequency of sequences with various mutation loads relative to baseline. (**C**) Frequency of lineages within various size bins relative to baseline. Here lineage size refers to the number of distinct sequences in a lineage.

In order to characterize the role of B-cell clonal proliferation in the replenishment of the repertoire, we assigned lineages to 3 bins depending on their degree of expansion (Fig. 6C). One can theorize two extreme modes of replenishment: in one scenario, the repertoire would get filled with a large diversity of clonally unrelated B-cells, none of which proliferate much; in the other scenario, the repertoire would be seeded by a single B-cell clone which proliferates to fill the entire repertoire. In reality, we observe an intermediate regime: all 3 lineage size bins contribute to replenishment (Fig. 6C). Patterns of clonal expansion are observed to be idiosyncratic to individuals: certain subjects display an increased importance of highly expanded lineages by the last time point (e.g. SR2 and SR6, also transiently the case for SR1) suggesting proliferation with somatic hypermutation (cf. Fig. 6B); another subject (SR3) displayed a markedly non-expanded, naïve phenotype at the end of depletion with mostly IgD/IgM antibodies showing few mutations (Fig. 6A,B,C); in yet another group of subjects, the importance of the 3 lineage size bins relative to each other at the last time point was comparable to baseline (SR1, SR5 in Fig. 6C).

## Discussion

We have characterized the role of B-cells in SSc-PAH at the level of the B-cell receptor. We identified an anomaly in the process of VDJ recombination, namely underuse of the IGHV2-5 segment (particularly in the isotype-unswitched compartment, Fig. 2F) and noted an interesting bimodal effect at the extremes of peripheral B-cell development: SSc-PAH participants displayed elevated proportions of both naïve BCRs (Fig. 2A) and of highly secreted antigen-experienced BCRs (Fig. 2B,C,D). It is worth noting that the presence of any infection at baseline would have made subjects ineligible to participate in the present study, so that any excess proportion of plasmablasts cannot be the result of current infection. Instead, the excess may suggest a sustained immune response against self-antigens, a hypothesis which is further corroborated by the observation of an increased proportion of near-fixed somatic mutations likely to arise from clonal selection (Fig. 2E). Certain of these disease signatures are clearly transiently reversed by CD20^+^ B-cell depletion, namely the IGHV2-5 gene usage (Fig. 3C) and the naïve proportion (Fig. 3A,B).

The observation of B-cell depletion in one group of study subjects (Fig. 1) allowed us to measure rates governing BCR diversity generation and dissipation in humans *in vivo* (Fig. 4A,B), demonstrating the high frequency of apoptosis of naïve cells compared to activation, as well as the precedence of somatic mutation over class-switch recombination in activation. More importantly, these measurements, combined with trends relating replenishment rates and depletion durations to baseline information (Fig. 4C,D), constitute a proof of principle for the prediction of immunologic depletion and replenishment time courses from baseline information alone. We compared the reconstituted repertoire to the baseline repertoire and noted a remarkable resiliency of summary properties such as VDJ frequencies, mutation histograms and isotype frequencies (Fig. 5A), indicating that the diversity generation mechanisms of VDJ recombination, somatic hypermutation and class-switch recombination are intact after depletion and able to regenerate a normal B-cell repertoire in a timespan of months.

We also dissected the mechanism of B-cell replenishment after depletion. We found that release of antigen-experienced cells into the periphery – likely plasma cells released from the bone marrow – does (Fig. 5B) or does not (Fig. 5C) precede the onset of naïve repopulation depending on the individual. Antibody repertoire sequencing exposed the lineage structure of the recovering B-cell ensemble, revealing subject-to-subject heterogeneity in the pattern of clonal expansion after depletion (Fig. 6C).

## Materials and Methods

Protocols were approved by the Institutional Review Board at Stanford University and at all Autoimmunity Center of Excellence sites providing samples for this study; participants gave informed consent. IGH repertoires were obtained using previously published protocols (31, 32) with small modifications, via the following steps: RT with IGH constant domain specific primers containing random barcodes (Table S2) on 250 ng total RNA extracted from peripheral blood mononuclear cells (PBMCs); second-strand synthesis with IGH variable domain specific primers containing random barcodes (Table S3); Ampure purification (Beckman Coulter); PCR amplification incorporating Illumina adapters and sample multiplexing indexes; 150-bp paired-end NextSeq sequencing (Illumina); analysis using pRESTO (33), IMGT/HighV-QUEST (34), Change-O (35), and R scripts to be deposited at https://github.com/cdebourcy/scleroderma. For more technical descriptions, see Supporting Information (*Supplement on Antibody Repertoire Sequencing and Analysis*).

## Acknowledgments

The present paper resulted from an Autoimmunity Centers of Excellence study. The Autoimmunity Centers of Excellence (ACE) is a research consortium supported by the National Institute of Allergy and Infectious Disease (NIAID/NIH). We thank the ACE ASC01 study group for providing SSc-PAH samples; Sally Mackey for coordinating the clinical studies that provided the healthy subjects; Sue Swope and Michele Ugur for conducting study visits and collecting samples from healthy subjects; Xiaosong He for providing access to samples from healthy subjects; Holden Maecker, Jackie Bierre and Ben Varasteh (Stanford Human Immune Monitoring Core) for sample banking; Norma Neff and Gary Mantalas (Stanford Stem Cell Genome Center) for assistance with sequencing; and Lolita Penland, Derek Croote, Felix Horns and Jacob Glanville for discussions. The research on SSc-PAH subjects was supported by NIH Grants U19 AI046374 (to Michael Holers of the Denver Autoimmunity Center of Excellence) and P01 HL014985, R01 HL122887 (to M.R.N.). The research on healthy subjects was supported by NIH Grant U19 AI057229 (to M.M.D.). The clinical project on healthy subjects was supported by NIH/National Center for Research Resources Clinical and Translational Science Award UL1 RR025744. The ClinicalTrials.gov numbers from the studies from which samples were used are NCT01086540 (SSc-PAH cohort) and NCT02133781, NCT03020498, NCT03022396, NCT03022422 (healthy cohort). C.F.A.d.B. was supported by an International Fulbright Science and Technology Award and a Melvin & Joan Lane Stanford Graduate Fellowship.

## Supporting Information

### Supplement on Antibody Repertoire Sequencing and Analysis

#### IGH repertoire sequencing

The assay used in the present study proceeded in the following steps, which were a minor variation of published protocols (22, 31, 32): PBMC isolation from whole blood using a Ficoll gradient according to Stanford Human Immune Monitoring protocols (storage by freezing in 10% (vol/vol) DMSO/40% (vol/vol) FBS); total RNA extraction from thawed cells using the Qiagen AllPrep kit; RT on 250 ng total RNA using SuperScript III reverse transcriptase (Life Technologies) and IGH constant domain specific primers containing 8 or 12-nucleotide random barcodes synthesized by Integrated DNA Technologies (Table S2); second-strand synthesis using Phusion High-Fidelity DNA Polymerase (New England Biolabs, 98 °C for 4 min, 52 °C for 1 min, 72 °C for 5 min) and IGH variable domain specific primers containing 8 or 12-nucleotide random barcodes synthesized by Integrated DNA Technologies (Table S3); two rounds of cDNA purification with Ampure XP beads (Beckman Coulter) at a 0.8:1 ratio; PCR amplification using Platinum Taq DNA Polymerase High Fidelity (Life Technologies) and primers containing sample-multiplexing indexes combined with Illumina adapters; another round of purification with Ampure XP beads (0.7:1 ratio); sample pooling at equal volume; gel-purification using E-Gel EX Agarose Gels 2% (Invitrogen) and Freeze ‘N Squeeze DNA Gel Extraction Columns (Bio-Rad); 150-bp paired-end NextSeq sequencing (Illumina).

#### Data pre-processing

Data pre-processing was carried out in a manner analogous to a previous study (22). First, we carried out the following steps of the pRESTO (33) suite (version 0.4.8): raw read filtering with a threshold of 20 for the mean Phred quality; removal of PCR primer sequences from both ends in two passes, dealing first with the random barcodes of length 8 nucleotides and then with the random barcodes of length 12 nucleotides, annotating each read with its barcode; consolidation of read groups with identical barcodes into a consensus sequence after alignment based on primer sequences; removal of read groups above a 0.1 error threshold against the consensus sequence; removal of read groups below a 70% agreement on constant-domain primer identity; merging of paired-end mates into a single sequence based on the read group consensus; removal of pairs below overlap significance threshold p < 10^−5^ or above overlap error threshold 0.3; removal of sequences containing more than 10 ambiguous inner nucleotides,;assignment of isotypes and subisotypes based on constant-domain sequence match with removal of sequences above error threshold 0.2; trimming of the constant-domain sequence; assignment of molecular abundances (number of paired-end barcodes observed for the same IGH sequence); assignment of consensus counts (number of read pairs supporting an IGH sequence); removal of sequences with consensus count under 2.

Next: IMGT/HighV-QUEST (34) was used for VDJ gene assignment; non-functional sequences were removed; TIgGER (25) was used to determine V-genotype and correct V-allele calls (after pooling of time points for the longitudinal experiment, not applicable to the baseline comparison experiment); a priori indistinguishable alleles (based on the sequenced region) were collapsed; Change-O (version 0.3.3) (35) was used to infer a germline sequence for each observed sequence, masking the CDR3 region with ambiguous nucleotides; low-abundance sequences only 1 nucleotide away from a high-abundance sequence (abundance threshold: 20) with an identical combination of V-gene, J-gene and CDR3 length were removed as they might have resulted from RT errors; finally, sequences with identical combinations of V-gene, J-gene and CDR3 length were clustered into lineages using single-linkage clustering on the CDR3 nucleotide sequence with a Hamming distance cutoff of 0.1 times the CDR3 length (using the R package ‘stringdist’ (36)). Snakemake (37) was used to manage computational workflows.

#### UniFrac calculation

UniFrac (21) distances were computed as described in a previous antibody repertoire study (22), subsampling each repertoire/time-point to a depth of 10^4^ distinct sequences if the total number of available sequences exceeded that number.

#### Mutation analysis

Reported nucleotide mutation counts were based on mismatches between each observed sequence and its inferred germline sequence. Reported AA changes for the V-segment were extracted from IMGT/HighV-QUEST output.

#### Plots

For Fig. 2C, abundance histograms were generated with a bin size of 20 and the lowest bin (which was the dominant one) was omitted from the plot to allow the tail of the distribution to be visualized. In Fig. 2D,F, points straddling the lower edge of the panel correspond to the value 0, which cannot be rendered on the logarithmic scale.

#### Statistical tests

P-values are from two-sided Wilcoxon-Mann-Whitney tests except for the correlation tests (two-sided PPMCC tests). Significance codes in figures: “*” 0.01 ≤ p < 0.05, “**” 0.001 ≤ p < 0.01, “***” 0.0001 ≤ p < 0.001.

## Supporting Figures

**Fig. S1.**
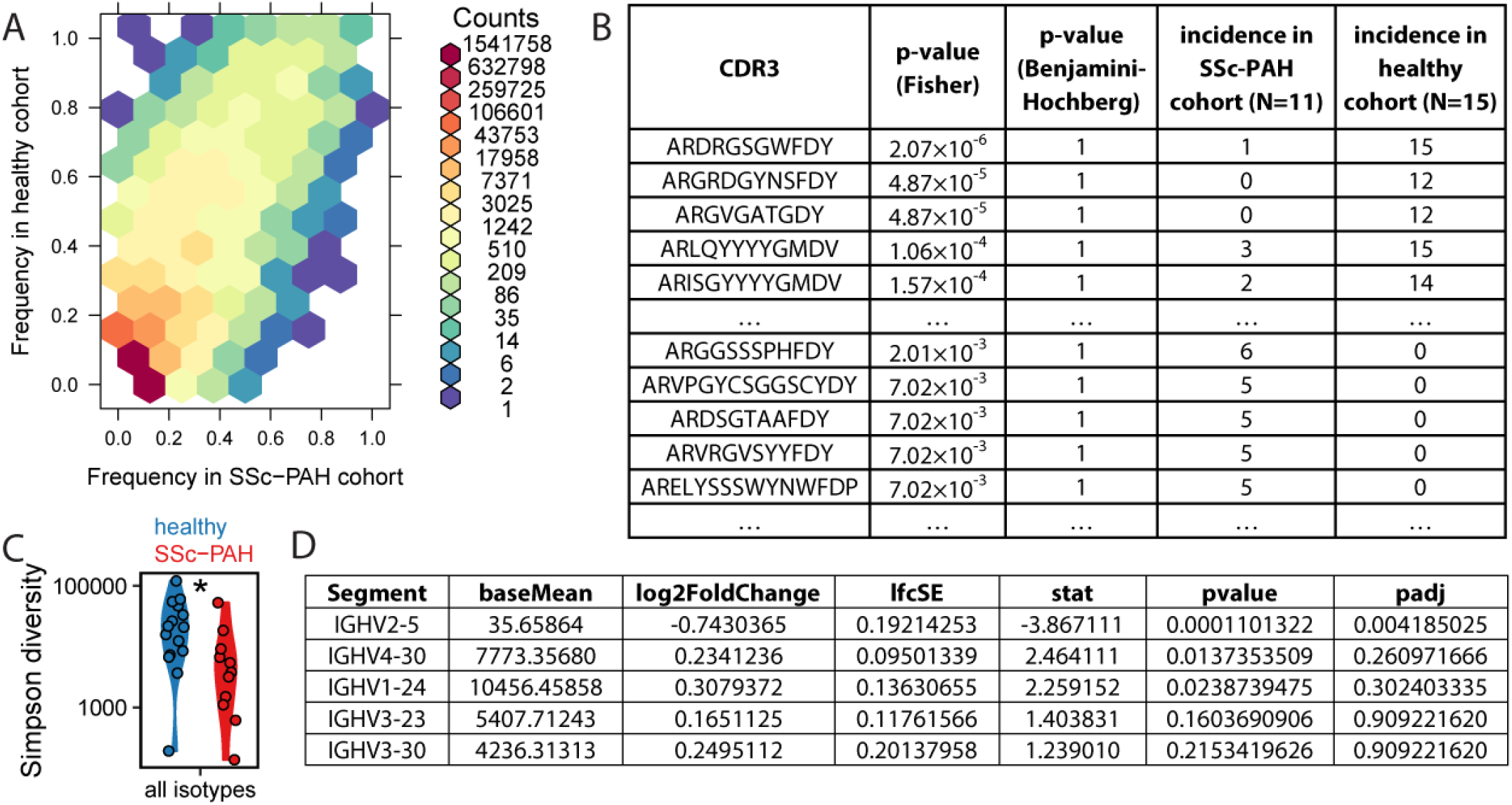
Comparison of healthy and SSc-PAH cohorts at baseline. (**A**) Two-dimensional histogram of AA CDR3 sequences indicating prevalence in the two cohorts. For each distinct AA CDR3 sequence, we determined which subjects contained that sequence or any of its one-mismatch derivatives in their repertoire. The frequency of the sequence in a cohort is the fraction of subjects in the cohort that contained at least one relevant transcript. It is observed that most sequences are low-prevalence, and that prevalences in the two cohorts are generally similar with no obvious outliers. (**B**) The top part of the table shows the five AA CDR3 sequences with the overall lowest unadjusted p-values from Fisher’s Exact Test. The bottom part shows five sequences with the lowest unadjusted p-values that were more prevalent in the SSc-PAH cohort than the healthy cohort. (**C**) Simpson diversity of the overall repertoire (1 divided by the sum of the squared fractional abundances of all sequences; shown on a logarithmic axis). Each point corresponds to one subject. (**D**) Differential gene expression analysis results output by DESeq2 (38), contrasting total repertoires for the SSc-PAH cohort relative the healthy cohort. “padj” is the Benjamini-Hochberg adjusted p-value. Only the 5 genes with the lowest padj-value are displayed; only IGHV2-5 was statistically significant.

**Fig. S2.**
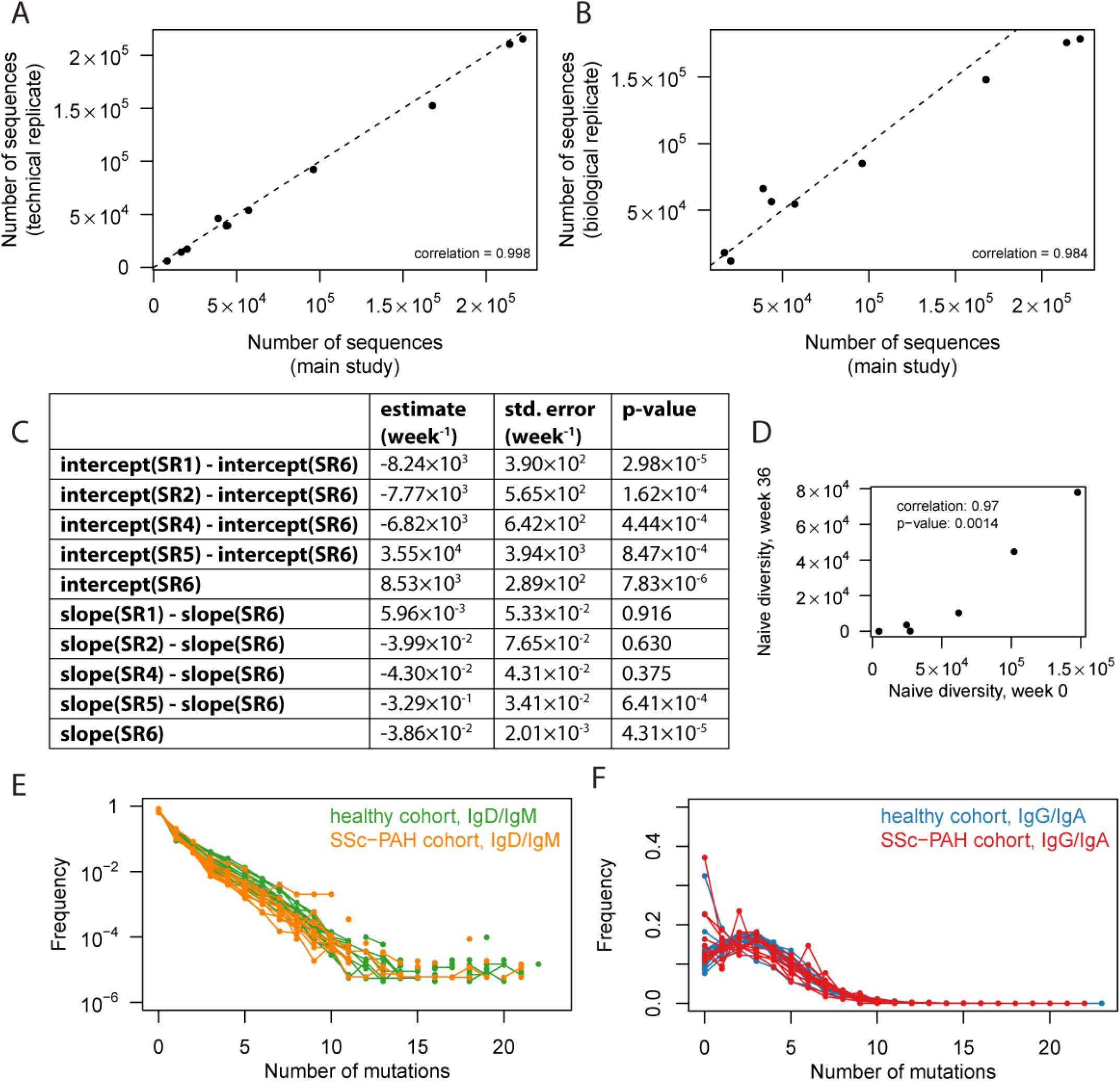
Calculation of rates governing repertoire homeostasis. (**A**) Observed sequence diversity is consistent across technical replicates. Values displayed on the horizontal axis correspond to the SSc-PAH baseline data from the longitudinal experiment described in the main text; technical replicates were generated by resequencing the corresponding libraries at a comparable depth. (**B**) Observed sequence diversity is consistent across biological replicates. Values displayed on the horizontal axis correspond to the SSc-PAH baseline data from the longitudinal experiment described in the main text; biological replicates were generated by making new libraries from different RNA aliquots of the same samples and sequencing them at a comparable depth (same data as the SSc-PAH group of the cross-sectional experiment described in Fig. 2 of the main text). The same mass of input RNA (250 ng) was used to make all libraries; samples for which insufficient RNA was left over from the main study were omitted (SR3 baseline, SP5 baseline). (**C**) Fit coefficients for a linear model of the form 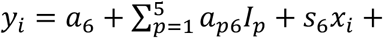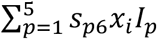 with standard errors and p-values for rejecting the null hypothesis that a coefficient is zero. Here (*x_i_*,*y_i_* is an 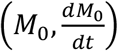 data point; *I_p_* is an indicator value with value 1 if the sample (*x_i_*,*y_i_*) is from participant *p* and 0 otherwise; *a_p6_* = *a_p_* − *a_6_* and *s_p6_* = *s_p_* − *s_6_* where *a_p_* and *s_p_* are the intercept and slope in the linear model applicable to the 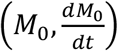 data of participant “SR”p (p=1,2,4,5,6). M_0_ trajectories were only modeled from the week 24 visit onwards to ensure onset of replenishment had already occurred; for participant SR1 it was necessary to begin later at week 36 for the trajectory to be monotonically increasing. Before conducting the fit, M_0_ temporal trajectories were first lightly smoothed (each 2 successive values were averaged and assigned to the middle of the 2 corresponding time points); then *dM_0_/dt* values were approximated as 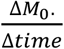 from each pair of adjacent time points and the average M_0_ of the same two time points was used as the corresponding M_0_-value to give an 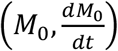 pair. Participant SR3 is omitted because an insufficient number of time points after onset of depletion were available; SR6 was chosen as the reference for slope and intercept because SR6 was tracked over the longest period of time. The table spells out “intercept” and “slope” instead of *a* and *s*, e.g. intercept(SR5) instead of *a_5_* and slope(SR5) instead of *s_5_*. Note that slope(SR1) – slope(SR6), slope(SR2) – slope(SR6) and slope(SR4) – slope(SR6) were all indistinguishable from zero (p-values above 0.1), suggesting that the slope of the linear model is similar for 4 out of the 5 modeled participants. (**D**) Number of non-mutated IgM or IgD sequences at baseline versus week 36. (**E**) Distribution of mutation loads in the IgD/IgM compartment at baseline. The distribution roughly corresponds to an exponential decay. (**F**) Distribution of mutation loads in the isotype-switched compartment at baseline. The distribution tends to be peaked at a non-zero value.

**Fig. S3.**
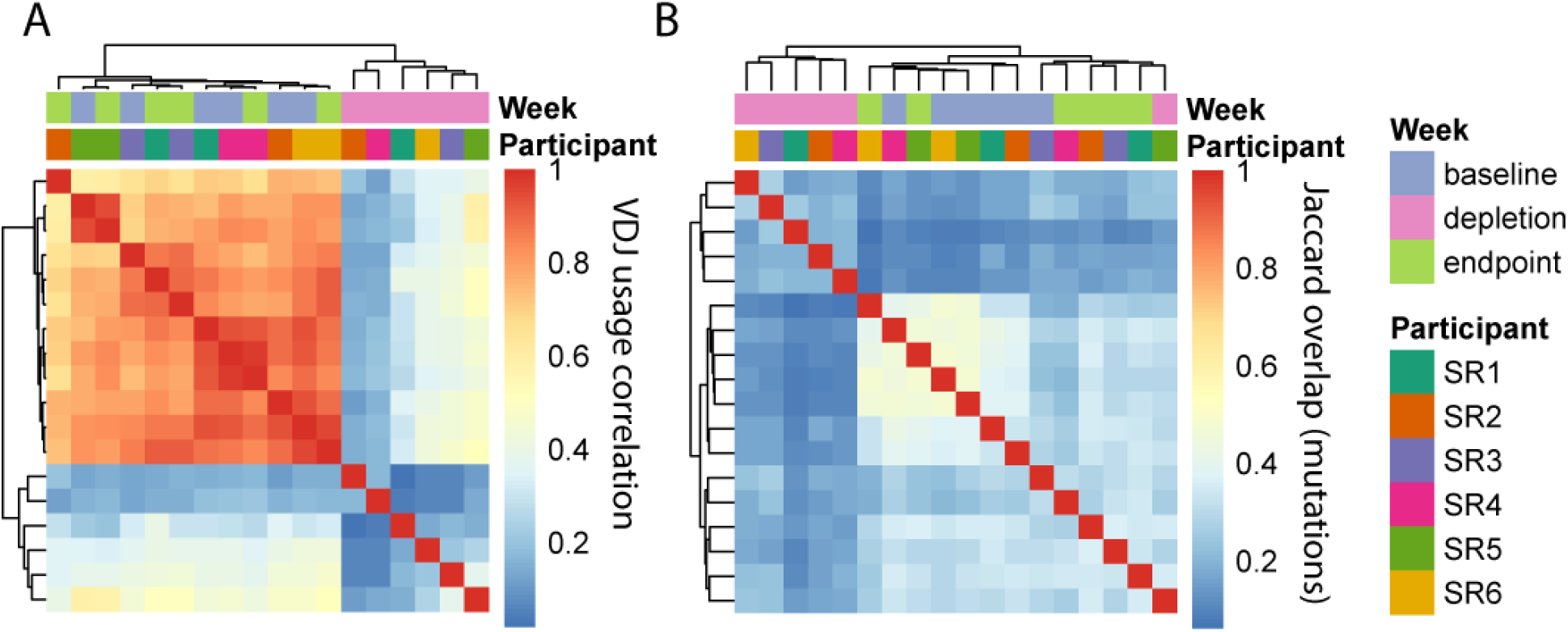
Assessment of repertoire elasticity under depletion. (**A**) Comparison of VDJ usages (sequence-weighted) between participants and time points, displayed as a correlation heatmap. “Depletion” corresponds to the week 4 sample, “endpoint” to the last time point sampled. Rows and columns were ordered using complete-linkage hierarchical clustering with Euclidean distance. (**B**) Comparison of AA mutation sets between participants and time points, displayed as a heatmap of Jaccard overlap indices. Jaccard indices of the sets of position-labeled AA changes were computed separately for each V-gene and then averaged over all V-genes. “Depletion” corresponds to the week 4 sample, “endpoint” to the last time point sampled. Rows and columns were ordered using complete-linkage hierarchical clustering with Euclidean distance.

**Table S1.**
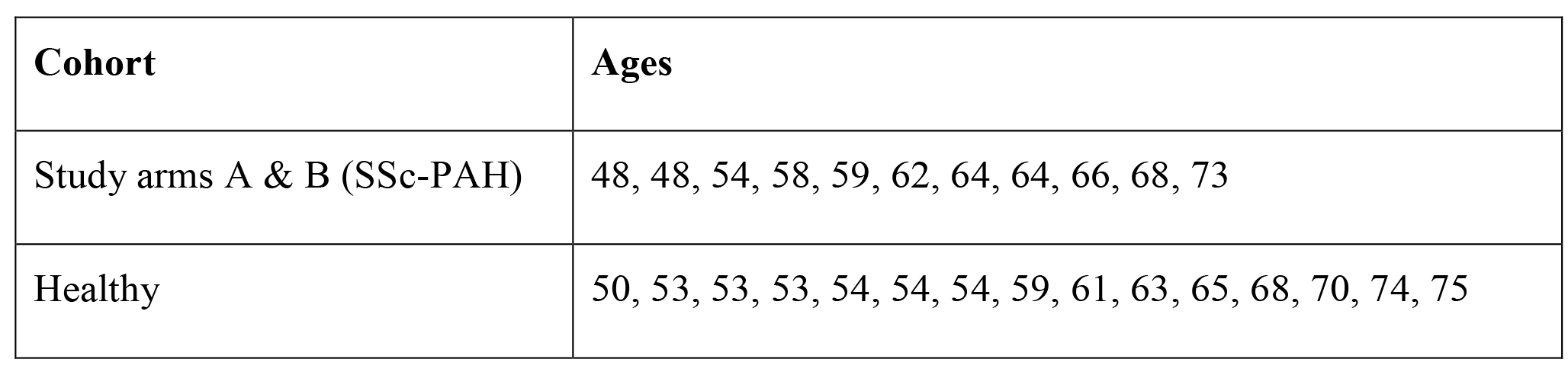
Aggregate demographic information on study participants.

**Table S2.**
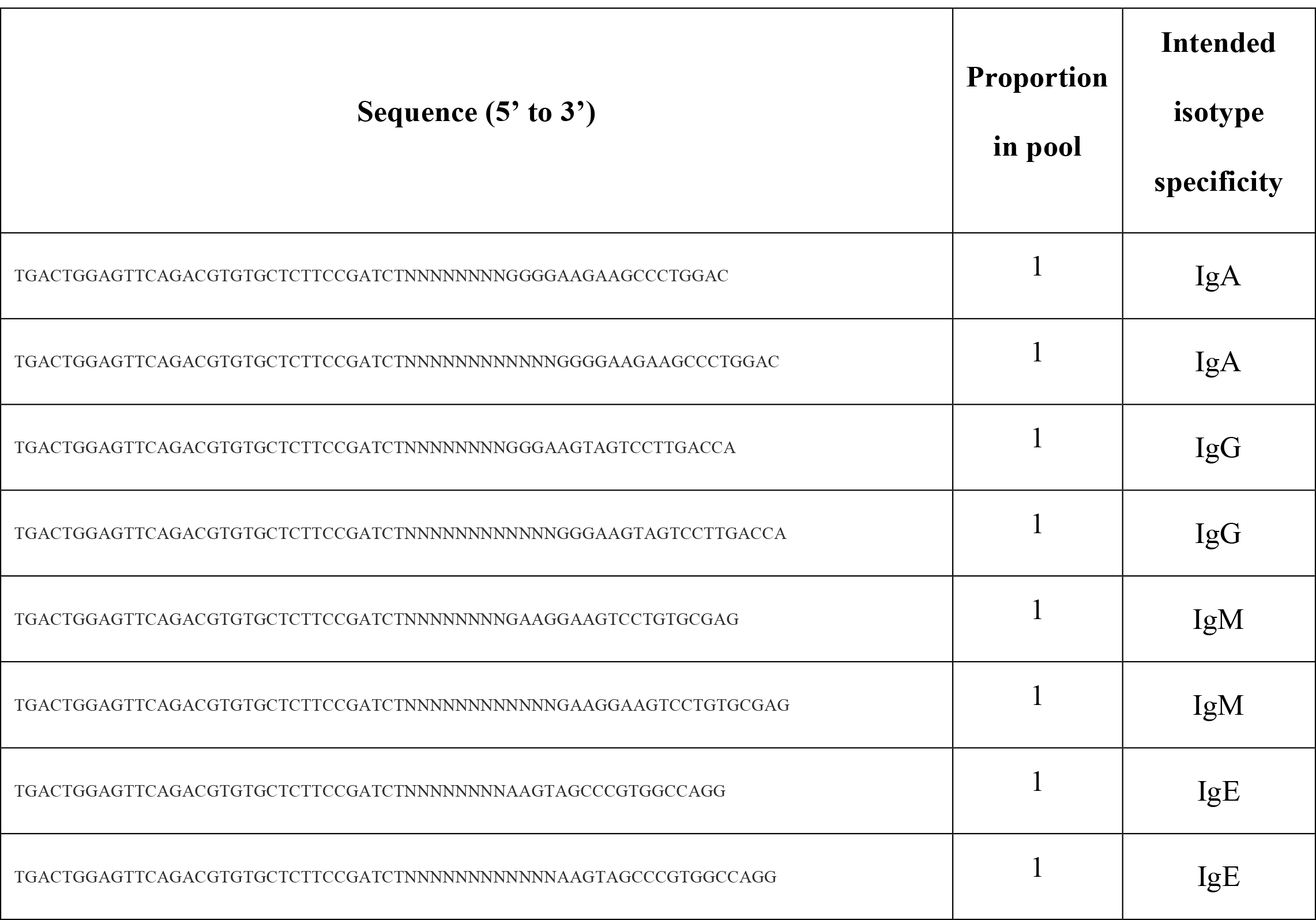
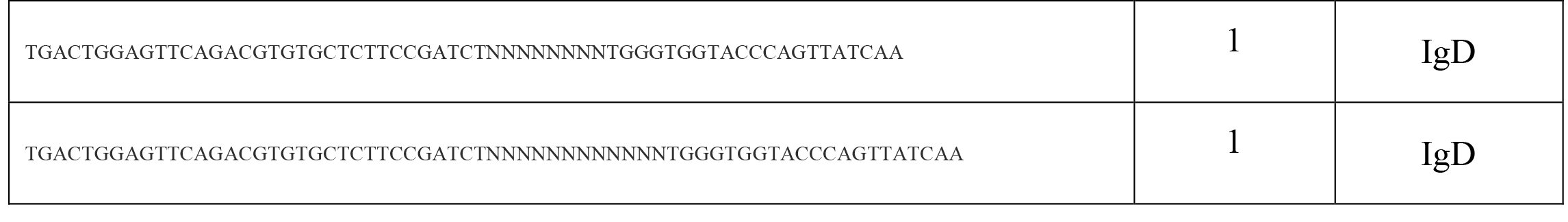
IGH constant-region primer pool. Follows a previously published protocol (32). Ns correspond to random nucleotides, to be used as a barcode for recognizing unique molecules in the starting RNA material.

**Table S3.**
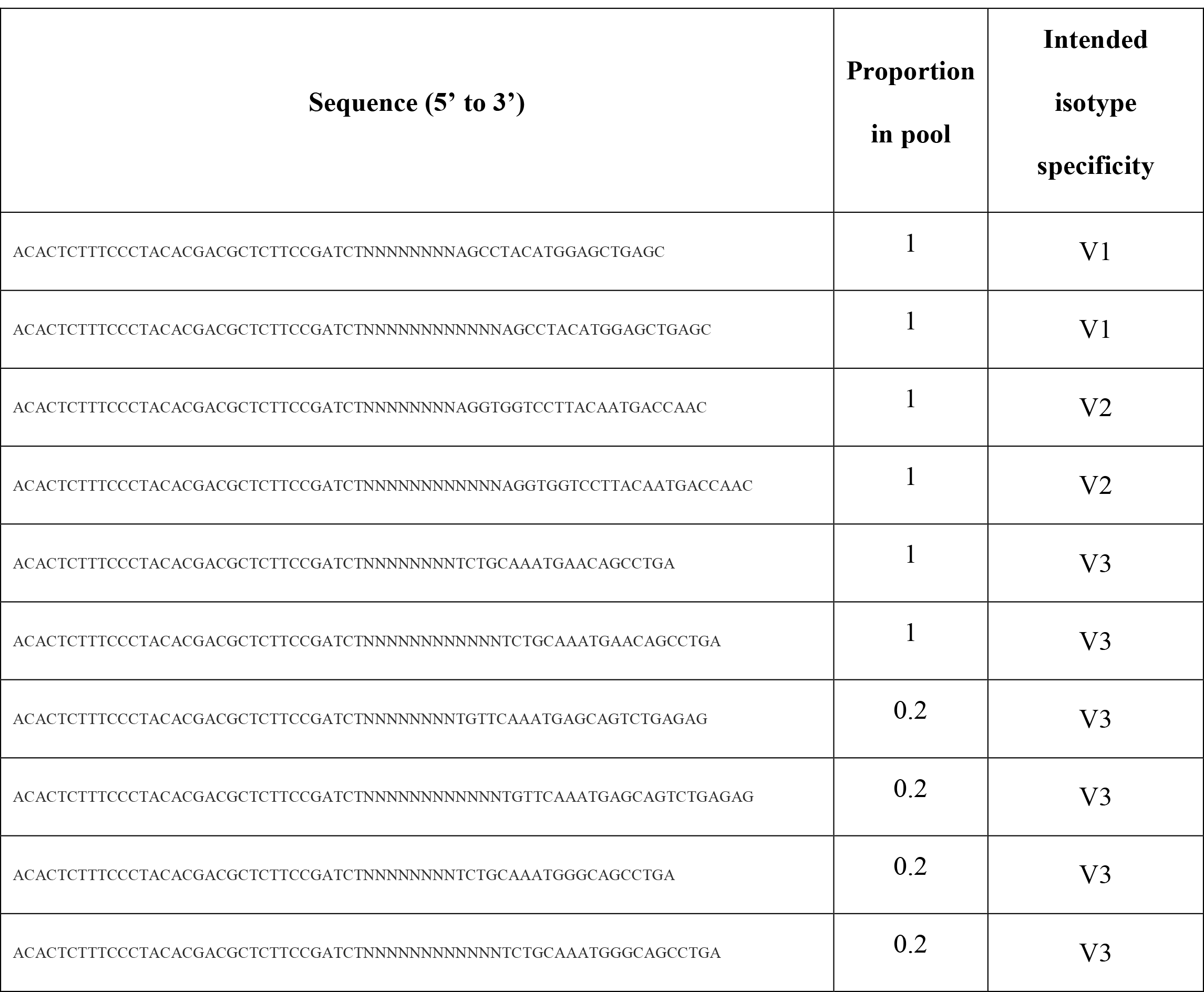
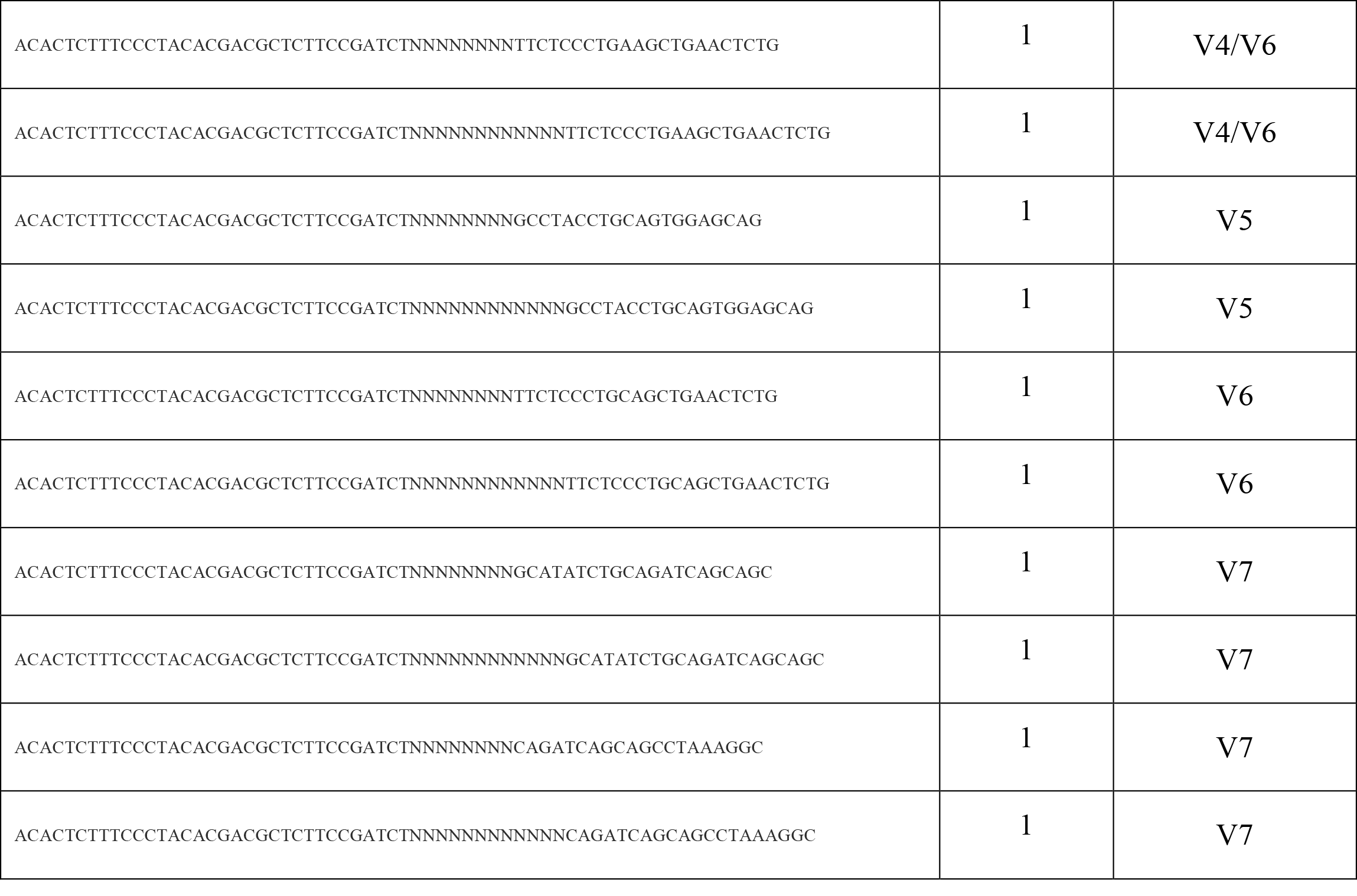
IGH variable-region primer pool. Based on a previously published protocol (31). Ns correspond to random nucleotides, to be used as a barcode for recognizing unique molecules in the starting RNA material.

